# Evidence Of We-Representation In Monkeys When Acting Together

**DOI:** 10.1101/2021.05.14.444171

**Authors:** Irene Lacal, Lucy Babicola, Roberto Caminiti, Simone Ferrari-Toniolo, Andrea Schito, Laura E. Nalbant, Rajnish K. Gupta, Alexandra Battaglia-Mayer

**Author notes:** Department of Physiology, Development and Neuroscience, University of Cambridge, Cambridge, CB2 3EG, UK.

## Abstract

A hallmark of successful evolution resides in the ability to adapt our actions to those of others, optimizing collective behaviour, to achieve goals otherwise unattainable by individuals acting alone. We have previously shown that macaques constitute a good model to analyse joint behavior, since they are able to coordinate their actions in a dyadic context. In the present work, we investigated whether monkeys could improve their joint-action performance, under special visuomotor conditions. The behavior of 5 monkeys was analyzed in isometric center-out tasks, requiring hand force application in different directions, either individually or together with a partner. Manipulating the presence or absence of a pre-instruction about the future action condition (SOLO or TOGETHER), allowed us to investigate on the existence of a “we-representation” in macaque monkeys. We found that pre-cueing the future action context increased the chances of dyadic success, also thanks to the emergence of an optimal kinematic setting, that ultimately facilitates inter-individual motor coordination. Our results offer empirical evidence in macaques of a “We-representation” during collective behavior, that once is cued in advance has an overall beneficial effect on joint performance.

## INTRODUCTION

The ability to coordinate actions among different individuals, in order to achieve goals unattainable by single subjects acting alone, has been observed across species. It is an expression of collective behavior, maximally developed in humans and considered as a key-element of successful evolution of our species (Boyd, 2018). Although several forms of apparent coordination have been observed even for very simple organisms such as insects, viruses and bacteria, our capabilities to coordinate with others is undoubtedly much more developed. A distinguishing element of such skills resides in our special ability to predict others’ actions and to flexibly adjust the own behavior to that of our co-agent (Boyd, 2018). A particular form of collective coordination is expressed during motor interactions, through inter-individual motor coordination. The terms “joint action” and “motor coordination”, despite referring often to similar contexts during social behavior across species, cannot be used as synonyms. Starting from the definition suggested by Sebanz et al. (2006), here we refer to the term ‘joint action’ as any form of motor interaction whereby two or more individuals, sharing an intention, coordinate their actions in space and time to achieve a common goal, while ‘motor-coordination’ describes the mere presence of a successful degree of synchronicity/complementarity among individuals, not necessarily requiring the assumption of a shared goal (Butterfill, 2017).

One of the first theoretical readout of social interactions in the context of motor control came from Wolpert et al. (2003) computational approach. From this perspective, socially connoted actions rely on similar mechanisms of individual actions. and would be based on the predictions of the consequences of our motor intentions, when interacting with the external environment. In social contexts, these predictions concern the effects of interactions of our actions with those of other individuals, to achieve a state estimation of the observed motor system. Therefore, engaging motor processes for action understanding is regarded as an efficient way for implementing the computations necessary for successful social interactions. These aspects go beyond the computational aspects of motor interactions, extending into the field of the psychology of collective behavior. During joint-action planning, a debated issue is whether motor preparation consists of separate predictive representations for one’s own and partner’s performance or whether it is grounded on a predictive action representation of the dyadic behavior. In the latter case the operating unit is the dyad, whose respective action representation would be the core of the so-called “We-representation” (Knoblich et al., 2011; Vesper et al., 2010) or “we-mode” (Gallotti & Frith, 2013). Under this scenario, when individuals join their forces to achieve a common goal, there is a *a priori* sense of doing something together, and not “on my own” and independently from the others. Therefore, joint performances would be guided by the collective goals, which are specified through dedicated motor representation (Della Gatta et al, 2017) rather than by each individual contribution.

In our previous work, we have shown that macaques are able to coordinate their action to achieve a common goal, by modulating their behavior on the basis of new task demands imposed by the dyadic context (Visco-Comandini et al. 2015; Ferrari-Toniolo et al. 2019). We observed that interacting comes with a cost, which affects performance, not only in non-human primates, but also in humans, whose joint-action abilities emerge during childhood (Satta et al. 2017). At the neural level, we provided the first evidence of a motor representation of joint-action (Ferrari-Toniolo et al. 2019) in population of neurons (‘joint-action cells’), that preferentially modulated their activity when a given action was performed in a dyadic context.

In the current work, we studied the influence on the joint performance, of pre-cuing the information about the action context (individual vs dyadic). Analyzing the macaque joint-action behavior in presence or absence of a pre-instruction, we searched for the existence of a putative “We-representation”, that similarly to that hypothesized in humans, may facilitate joint performace (Kourtis et al. 2019, Della Gatta et al 2017; Sacheli et al. 2018), by shaping action intentions at group level. To this aim, three couples of monkeys were trained to perform an isometric task, consisting in guiding a visual cursor on a screen from a central position to one of eight possible peripheral targets, by applying hand forces on an isometric joystick, either individually (SOLO) or in coordination with the partner (TOGETHER). In the latter condition, the animals had to coordinate their forces in direction and in time to achieve their common goal and to get their reward, as in Visco-Comadini et al (2015). The directional array augmented the level of spatial uncertainty, increasing across trials the re-adaptation demand to coordinate the own forces with a partner, given the kinematics variability during force application across directions. This allowed the analysis of the behavior under a wider range of motor scenarios. Importantly, two different tasks were adopted to instruct the monkeys about the type of action to be executed (SOLO vs TOGETHER). In the first (no-Pre-Instructed task, noPI), the social cue and the direction of the force to be applied were provided simultaneously. In the Pre-Instructed task (PI), instead, the “social condition” was pre-cued and then the information about the force direction followed. The serial order task structure allowed the animals to be prepared to the type of action first, and only subsequently to shape a motor plan for the force direction.

Manipulating the presence/absence of a pre-cue about action contexts provides a useful tool to highlight the existence of a “We-representation”. The aim was to provide an experimental evidence of the animals’ sense to act together, by disentangling the planning processes associated to joint action from those related to the actions per se. Our hypothesis is that if the animals lack the sense of “we-ness” (Gallotti & Frith, 2013), a pre-cue provided about action context (SOLO vs TOGETHER) should not influence joint behavior, since an identical motor representation would be adopted by the animals, acting alone or together. On the contrary, if joint action plans rest on a predictive representation of the collective action, and therefore a “we-representation” is available, pre-cuing the type of representation (I vs we) of future action should benefit joint performance.

## METHODS

### Animals

Five male rhesus monkeys (Macaca mulatta) were used for this experiment (**Fig. 1B**: Monkey S, 6.5 Kg; Monkey K, 7.5 Kg; Monkey C, 9 kg; Monkey D, 8.5 Kg, 11.5 Kg (D*); Monkey P, 11 Kg). D* indicates data obtained from Monkey D tested around one year later. The five animals were paired to form three couples, as follows: Monkeys S and K (couple SK); Monkey C and D (couple CD); Monkey D* and P (couple DP). All efforts were made to optimize animal welfare. Animal care and housing procedures were in conformity with European (Directive 63-2010 EU) and Italian (DL. 26/2014) laws on the use of non-human primates in scientific research.

**Fig. 1.**
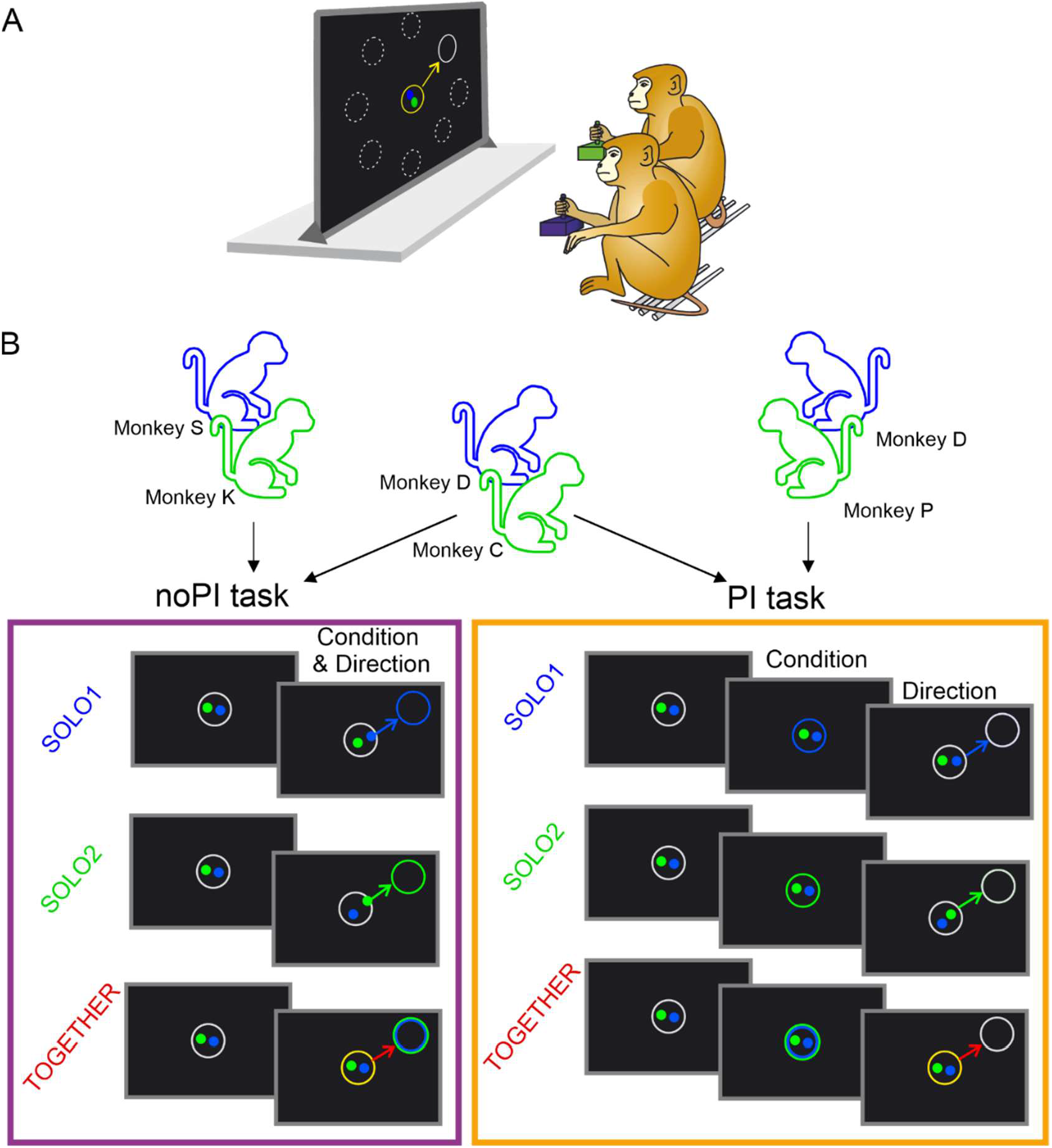
Experimental apparatus and tasks. **A.** Two monkeys seating side-to-side in front of a monitor performed an isometric directional center-out task consisting in controlling a colored visual cursor (e.g. blue, Monkey 1; green, Monkey 2) from a central position to a peripheral target, under two different conditions (SOLO or TOGETHER). **B.** Five animals (Monkeys S, K, C, D and P) were paired to form three couples (SK, CD, DP) of co-acting partners. The animals were required to guide their own cursor from the center to the peripheral target in order to receive their reward dose, either individually (SOLO trials) or jointly with their partner (TOGETHER trials). In the latter case, the monkeys had to move a common visual object (yellow circle) from the center toward a peripheral target, by coordinating their hand forces to maintain an intercursors distance <5 DVA for the entire duration of cursors’ motion. Once the yellow circle overlapped the peripheral target and kept in the new position for a variable THT (100-200ms), both animals received their liquid reward dose. Two different task structure (noPI and PI task) were adopted, differing in the way the animals were instructed about the future action types. In the noPI task, the condition (SOLO1, SOLO2 or TOGETHER) and the direction of movement were instructed simultaneously (“Condition & Direction”). In the PI task, instead, trial type (“Condition”) was pre-instructed during a dedicated time interval lasting 700-1000 ms, after which the peripheral target location (“Direction”) was shown. SK and DP couple performed the noPI and PI task, respectively; while CD couple was tested in separated sessions, both in the noPI and the PI task.

### Experimental apparatus

During the experiment, two monkeys seated side-to-side on two primate chairs in front of a 40-inch monitor (100 Hz, 800-600 resolution, 32-bit color depth; monitor-eye distance: 150 cm; **Fig. 1A**). A security distance of 60 cm between animals was always guaranteed, in order to prevent physical contact. The experimental set-up was conceived to minimize, during data acquisition, any potential interaction outside the task, to avoid potential sources of uncontrolled variables. The orientation of the two chairs minimalized also visual contact. During the experiment, monkeys were required to use always the same arm, while the other was gently restrained. Each animal was trained to control a colored circular cursor (diameter: 0.6 degrees of visual angle; DVA) displaced on a black screen, by applying a hand force on an isometric joystick (ATI Industrial Automation, Apex NC) in two dimensions on the horizontal plane (sampling frequency: 1 kHz). The applied force was proportionally converted into a motion of the cursors on the x and y axis of the vertical plane of the monitor. The NIH-funded software package REX was used to control stimuli presentation and to collect behavioral events and force data.

### Behavioral task

All monkeys were required to perform an isometric directional center-out task consisting in moving a visual cursor from a central position to a peripheral target, in two intermingled conditions (SOLO or TOGETHER, see below for details about action conditions). The peripheral target was located in one of eight possible locations, at an eccentricity of 8 DVA.

In the SOLO condition, the animals were required to move their own cursor within the peripheral target, by applying a dynamic hand force on their own joystick. In the TOGETHER condition both monkeys had to move their cursors towards the peripheral target in a coordinated fashion, that is through an inter-individual coordination of their hand forces in speed, intensity and direction, in order to move a common visual object (represented by a yellow annulus) from the center to the peripheral location. The annulus was placed in the midpoint of the two cursors controlled by each animal. To instruct the monkey with an appropriate ‘social cue’ about the type of action (SOLO vs TOGETHER) to be executed, and/or about the final location to be reached by the controlled cursor, we used two different task structures: i) a Non Pre-Instructed task (noPI), where in each trial the information about the action type and force direction were provided all at once in a reaction-time paradigm; ii) a Pre-Instructed task (PI) where the two information were provided separately and sequentially: the action condition was first pre-instructed during a dedicated variable time interval, after which the directional cue was presented. The two task versions were presented in separate sessions.

The animal couples were assigned to the different type of task as follows: SK couple performed the noPI task (SK_noPI_), DP performed the PI task (DP_PI_), while CD couple performed both the noPI and the PI tasks, in the sequence CD_noPI_ and then CD_PI_.

Both noPI and PI tasks begun identically **(Fig. 1B)**, with the presentation of an outlined white circle (2 DVA in diameter) on the center of the screen, and each animal had to bring its cursor inside it and hold it there for a variable control time (CHT, 500-1000ms).

In the noPI task **(Fig 1B,** left), at the end of the CHT, a peripheral target (outlined circle, 2 DVA in diameter) was presented in one of eight possible positions (at 45° angular intervals) at an eccentricity of 8 DVA. Its color instructed the animals about the type of action (SOLO or TOGETHER condition) required to obtain a liquid reward. Therefore, in the noPI task, the social cute about action condition and future force direction were simultaneously instructed. Following the appearance of the peripheral target (GO signal) the animals were required to bring their own cursors (individually or together, see below for details), on the new location to get the reward, within a certain RT (100-800 ms limits).

In the PI task **(Fig 1B,** right), instead, the CHT was followed by a color change of the central target, which cued the monkeys on the future action condition (SOLO or TOGETHER) to be performed for a pre-instruction time (PIT, 1000-1500ms. During the PIT the animals were required to maintain their cursors within the central target, and when a white peripheral target (GO signal) appeared in one of the eight possible positions, they had to bring together or individually their own cursor toward the peripheral location within a given RT (100-800ms), as instructed by the color of the central target during the PIT. Cursors’ and instructing targets’ colors were always univocal for each monkey. In each dyad, monkey 1 was associated to the blue color and monkey 2 to green, which coincided to the colors of the respective cursors controlled by each animal. The SOLO condition for each monkey was thus instructed by blue or green target, while the TOGETHER condition was instructed by a bi-colored (blue and green) circumference. In the SOLO1/SOLO2 condition, a blue/green circle was presented, and Monkey 1/2 was instructed to move its own cursor within the peripheral target, by applying a dynamic hand force on the joystick. It was then required to maintain the cursor on that position for a variable target holding time (THT: 100-200ms), as to gain a reward (0.5 ml of fruit juice). During SOLO1 and SOLO2 trials, instead each partner was required to keep holding its cursor inside the central target for the entire duration of the trial, to gain 50% of the same liquid treat, irrespective of the partner’s action outcome (success or error). In the TOGETHER condition, when the peripheral target was presented, both monkeys had to move their cursors towards it in a coordinated fashion, to gain both the same amount of reward provided for a successful SOLO action (0.5 ml), after holding the final position for a variable THT (100-200ms). For the entire duration of their cursors’ motion, monkeys were constrained to stay within a maximum inter-cursor distance (ICD_max_) of 5 DVA, which was marked by a yellow circle encompassing the two cursors. The instantaneous position of the center of the yellow circle coincided with the mean value of the x and y coordinates of the two cursors. An ICD > ICD_max_ resulted in an “ICD error”, with the consequence of the trial abortion and none of the two agents obtained the reward. Within each session, trials corresponding to different action conditions (SOLO1, SOLO2, TOGETHER) and different directions were presented in an intermingled fashion and pseudorandomized within a session of minimum 192 successful trials (3 conditions, 8 directions, 8 replications). The number of sessions collected was different for each pair. For SK and for DP pairs, 72 and 41 sessions were collected, respectively. For CD couple, instead, 12 sessions for the noPI task, and 13 sessions for the PI task were acquired.

### Data analysis

#### Behavioral parameters

##### Successes and errors

To allow an overall comparison between the two action conditions, for each couple and each session, as an index of performance we computed the success rate (SR) for SOLO and TOGETHER trials, separately and independently from the direction factor. The variation in the performance of TOGETHER trials, across task types (noPI, PI) was evaluated with respect to that of SOLO trials, by computing for each session the difference between the SR associated to SOLO and TOGETHER condition (SR_SOLO_-SR_TOGETHER_). For TOGETHER trials, the rate of the ICD error (ER_ICD_) was also calculated (see above for its definition) and used as a further measure of performance accuracy, during the joint performance.

##### Reaction time and cursor peak velocity

In each SOLO and TOGETHER trial, the reaction time (RT) was defined as the time elapsing from the presentation of peripheral target to the onset of the cursor’s motion, which corresponded to the onset of the dynamic force application. The onset time was defined as the time at which the cursor’s velocity for at least 90ms exceeded by three standard deviations (SD) the average velocity signal, measured in the interval spanning from 50ms before to 50ms after the presentation of the peripheral target. The peak velocity (PV) was estimated for each trial, as the maximum value of the cursor’s tangential velocity.

##### Inter-individual differences

As a measure of the behavioral differences between the two subjects of each dyad, during both the planning and execution phase of a given task, we computed the inter-individual differences for the hand reaction times (IID_RT_) and the velocity peaks (IID_PV_). These were calculated for both SOLO and TOGETHER trials. In particular, for each *j*-th session the indexes IID_RT_ (and similarly the IID_PV_) were calculated as follows:

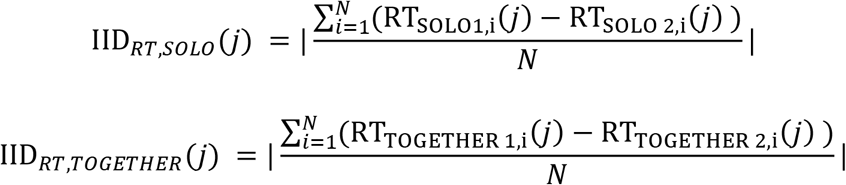

where *i* is the trial number and N is the total number of trials for the considered *j*-th session. For the SOLO condition, this index indicated the differences between the two subjects’ behavior when working independently from one another. In this case the differences were calculated after pairing the trials in random fashion. For the TOGETHER condition, the IID indexes have been considered as a measure of inter-individual motor coordination.

#### Statistical tests

Samples were first tested for normality though the Shapiro-Wilk test, due to its higher power compared to equivalent tests (Razali & Wah, 2011). Non-parametric statistical tests were applied when samples were not normally distributed. Wilcoxon rank sum test was used for two samples and Kruskal-Wallis test for k>2 groups, in both instances at a confidence level of a=0.05.

Success rates (SR) differences across trial types (SOLO1, SOLO2 and TOGETHER) were evaluated through Kruskal-Wallis test (factor: ‘trial type’, 3 levels), within each dataset. Dunn-Šidák test was used for multiple comparison between groups.

The difference in performance variation during the TOGETHER condition respect to SOLO one (SR_SOLO_-SR_TOGETHER_), evaluated in the noPI and PI tasks, was assessed through the Wilcoxon rank sum test either for each animal against all the other data sets through a post-hoc comparison or by pooling the data from all datasets.

To evaluate whether inter-individual differences in RTs and PVs correlate with the goodness of the TOGETHER performance, we adopted a Repeated Measure Correlation method (‘rmcorr’ package, R software, Bakdash & Marusich, 2018), which has the advantage to capture the across sessions relationship between the two variables that would be missed by using averaged data. IID_RT_/IID_PV_ and ER_IDC_ were calculated for each direction, within each session. In this analysis each session corresponded to one repetition. Therefore, a correlation was ultimately performed relative to a pool of 8 × *n* values, where *n* is the number of sessions for each dataset.

All statistics, but the Repeated Measure Correlation, were performed by use of MATLAB software (R2019a).

## RESULTS

### Performance across conditions and tasks

Five macaque monkeys, paired to form three couples, were successfully trained to perform isometric joint actions under different task conditions. In each experimental session, two animals seating side-to-side in front of a wide monitor (**Fig. 1**) had to guide each a visual cursor from the center to a peripheral target, by applying a force on their own isometric joystick. The isometric actions had to be required either in a SOLO fashion or jointly with the partner (TOGETHER), according to the instructions provided in each trial. In the SOLO condition, the animals were required to move their own cursor within the peripheral target individually, by applying a dynamic hand force on their own joystick. In the TOGETHER condition both monkeys had to move their cursors jointly towards the peripheral target in a coordinated fashion, that is through an inter-individual coordination of their hand forces in speed, intensity and direction, in order to move a common visual object (represented by a yellow annulus) from the center to the peripheral location (see Methods). The animals were instructed by an appropriate ‘social cue’ about the type of action be executed (SOLO vs TOGETHER), and /or by a visual cue about the final location to be reached by the controlled cursor. These instructions were provided in two different task structures: i) in the Non-Pre-Instructed task (noPI), the information about the action type and force direction (peripheral target to be hit by the moving cursor) were provided in each trial all at once in a reaction-time paradigm; ii) in a Pre-Instructed task (PI) the two information were provided separately and sequentially: the action condition (SOLO/TOGETHER) was first pre-instructed during a dedicated variable time interval, followed by the directional cue. The two task versions were presented in separate sessions.

We first evaluated the animals’ performance across action types (SOLO, TOGETHER) and different task structure, that is in absence (noPI) and presence (PI) of the pre-instruction about the type of action **(Fig. 1)**, by measuring the relative success rate (SR; **Fig. 2A**). We found, as previously shown (Visco-Comandini et al. 2015), that the SRs during the joint-action (TOGETHER) trials deviated from those of SOLO trials, being generally significantly lower (Dunn-Šidák test post-hoc comparison, p<0.05). The decrease of joint performance was stronger when the animals were exposed to the noPI task, namely when the monkeys were not pre-instructed about the future action type. The main goal of the present study was indeed to assess whether a pre-instruction of acting individually or jointly with the partner influenced the goodness of the overall performance. Thus, under the two different task contexts (PI vs noPI), we measured the deviation of the SR of TOGETHER trials from those of SOLO trials (SR_SOLO_-SR_TOGETHER_), by pooling the data from all datasets **(Fig. 2B)**. We opted to evaluate the joint-action SRs with respect to those of individual action, to avoid biases due to intrinsic variations in performances across single individuals. We found that, when the animals were prepared in advance to act together, the deviation diminished significantly (Wilcoxon rank-sum test, p = 1.67e-29, **Fig. 2B**), indicating that under such condition the inter-individual coordination improved dramatically by bringing the TOGETHER performance closer to the successful rates of the SOLO action. This phenomenon was observed in all five monkeys used in this study **(Fig. 2C;** Dunn-Sidák’s test, p<0.05, with the exception of D_C_ in PI not significantly different from S,K,D in noPI). The pre-instruction did not result to be similarly beneficial in the case of individual (SOLO) actions in all animals (Wilcoxon rank sum test, p>0.05; **Fig. 2D)**.

**Fig. 2.**
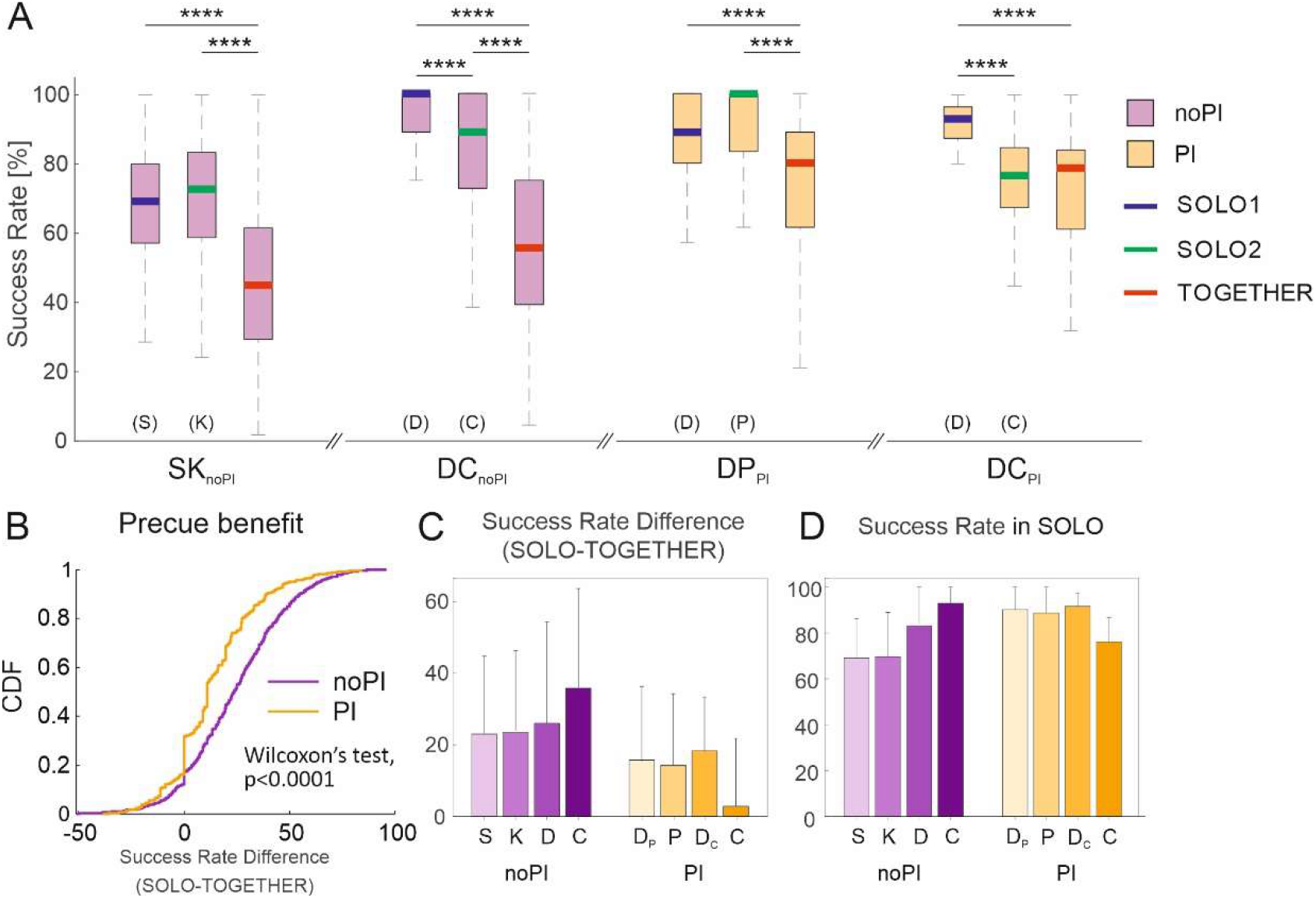
Task performance across SOLO/TOGETHER actions and in presence or absence of preinstruction. **A.** Box plots of success rates during SOLO1/ SOLO2 (median blue/green) and TOGETHER (median red) for each dataset, collected during the performance of noPI (purple boxes) and PI task (orange box). (****: Dunn-Šidák’s test, p<0.0001). **B.** Cumulative distributions of success rates differences between SOLO and TOGETHER trials. **C.** Mean differences between success rate during SOLO and TOGETHER trials, plotted for each animal tested in the noPI (purple) and PI (orange) task. **D.** Mean success rates of SOLO trials for each animal, tested in the noPI and PI task. Error bars in **C,D** indicate standard deviation.

### Effect of condition on monkeys’ kinematics

Given the facilitatory effect on dyadic performance of the anticipatory information to act in a joint context, we investigated on the nature of the influence that prior information exerts on monkeys’ joint behavior, so as to see which motor aspects were particularly affected, which in turn might foster inter-individual coordination. We focused our attention on the reaction times (RTs) and peak of velocities (PVs) of cursor motion, representative of the planning and execution phases of the isometric task, respectively. The modulation of these variables during joint performance relative to solo actions has already been documented in our previous study (Visco-Comandini et al. 2015). However, here we have re-evaluated the effect of the type of action on the kinematic profiles, separately for the noPI and PI sessions (**Fig. 3**). Our findings show that when pre-instructed about the future action condition (PI), in three cases out of four, a significant decrease of RTs in the TOGETHER condition with respect to the SOLO one **(Fig. 3A),** occurred. In the noPI task the results varied across animals **(Fig. 3A)**. Concerning the execution phase of the task, we found that during joint-action trials monkeys tended to slow down the speed of hand force application, as indicated by the decreased peaks of velocity (PV_SOLO_ > PV_TOGETHER_). This was generally true independently from the anticipatory instructions available to the animals.

**Fig. 3.**
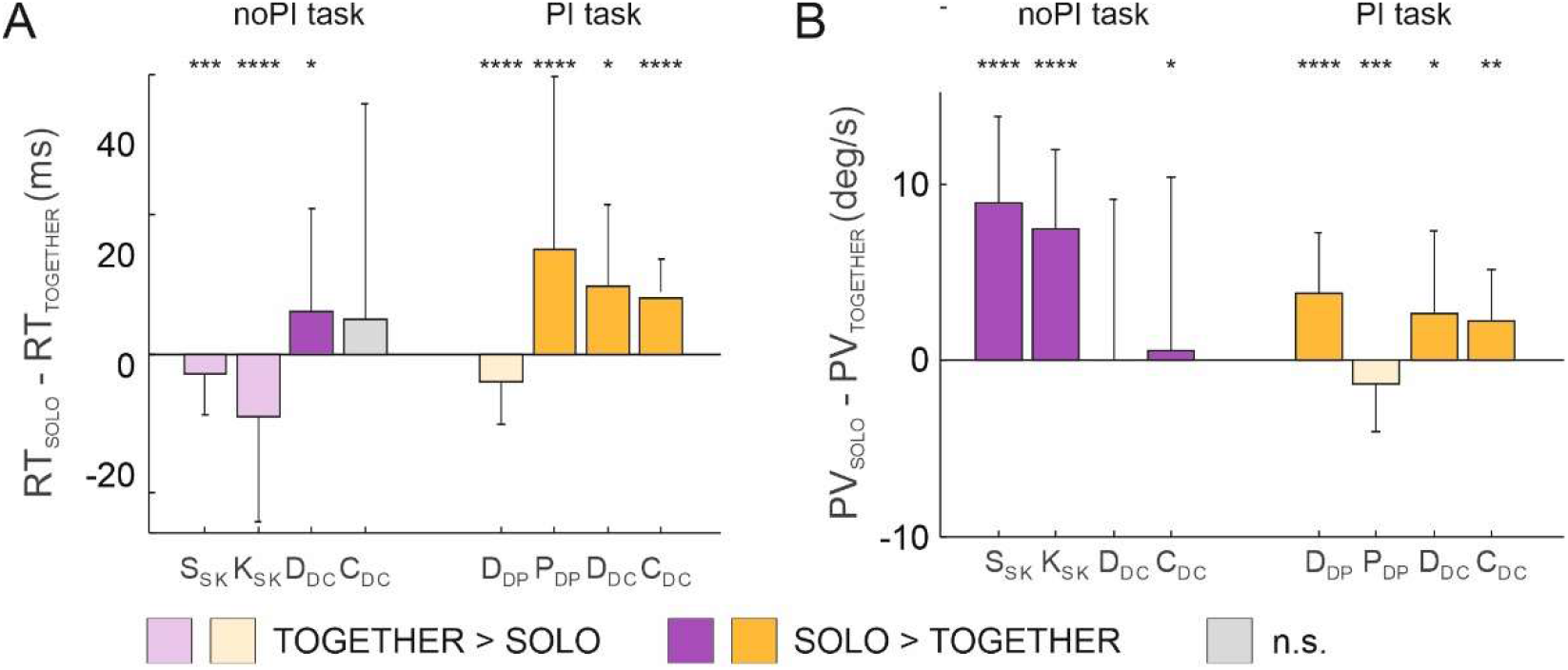
Effect of action condition on reaction times and velocity peaks. Difference between the median RTs (**A**) or median PVs (**B**) of SOLO and TOGETHER conditions, plotted for each animal tested during the performance of noPI (purple bars) and PI task (orange bars). Significant differences between SOLO and TOGETHER condition were evaluated through Wilcoxon rank sum test (*, p < 0.05; **, p < 0.01;***, p < 0.001; ****, p < 0.0001).

These findings suggest that there are behavioral invariances, such as a tendency to reduce the RTs in the joint performance, particularly in presence of a PI delay period, and to slow down the rate of force application, resulting in decreased cursor’s velocity, during the TOGETHER condition. However, beyond such general tendencies, we also observed inter-individual differences in the way each animal cope with the dyadic context. The next step is, therefore: (i) to understand if and how idiosyncratic differences between the two interacting partners can explain the emergence of variable strategies, (ii) how the entity of these differences can be influential on the goodness of joint performance, and (*iii*) whether the presence of the PI cue might be an influential factor in coping with these differences.

### Individual and inter-individual differences during SOLO action

We showed that the animals did not always modulate in a univocal fashion their behavior during joint performance, relative to their individual action. Hence, we asked whether the way each monkey responded to the demands imposed by the joint contexts depended on the degree of inter-individual difference between its own motor profiles and that of the partner. To this aim, we first measured to which extent in TOGETHER trials monkeys’ behavior differed from one another in individual behavioral parameters (i.e., RTs and PVs), as compared to SOLO trials **(Fig. 4)**. We found significant differences among animals’ RTs when acting individually, both in noPI and PI databases (Kruskal-Wallis’s test, noPI: χ2(3) = 3121.48, p = 0; PI: χ2(3) = 6383.59, p = 0; **Fig. 4A**). Similarly, also PVs differed across monkeys in both task types (noPI: χ2(3) = 1777.90, p = 0; PI: χ2(3) = 5282.21, p = 0; **Fig. 4B**). Post-hoc analysis confirmed that monkeys’ behavior differed in at least one of the two kinematic measure adopted in this study **(Fig. 4A-B**, bottom panels). We concluded that each monkey showed a spontaneous individual kinematic profile, which made its cursor’s motion diverse from that of its partner. Furthermore, we found that for each monkey both behavioral parameters significantly differed across directions (**Fig. 4A-B**, top panels). The question is whether the amount of divergence between the two “idiosyncratic” behavioral patterns, shown by the dyad’s members when acting alone, predicts their performance during joint-action.

**Fig. 4.**
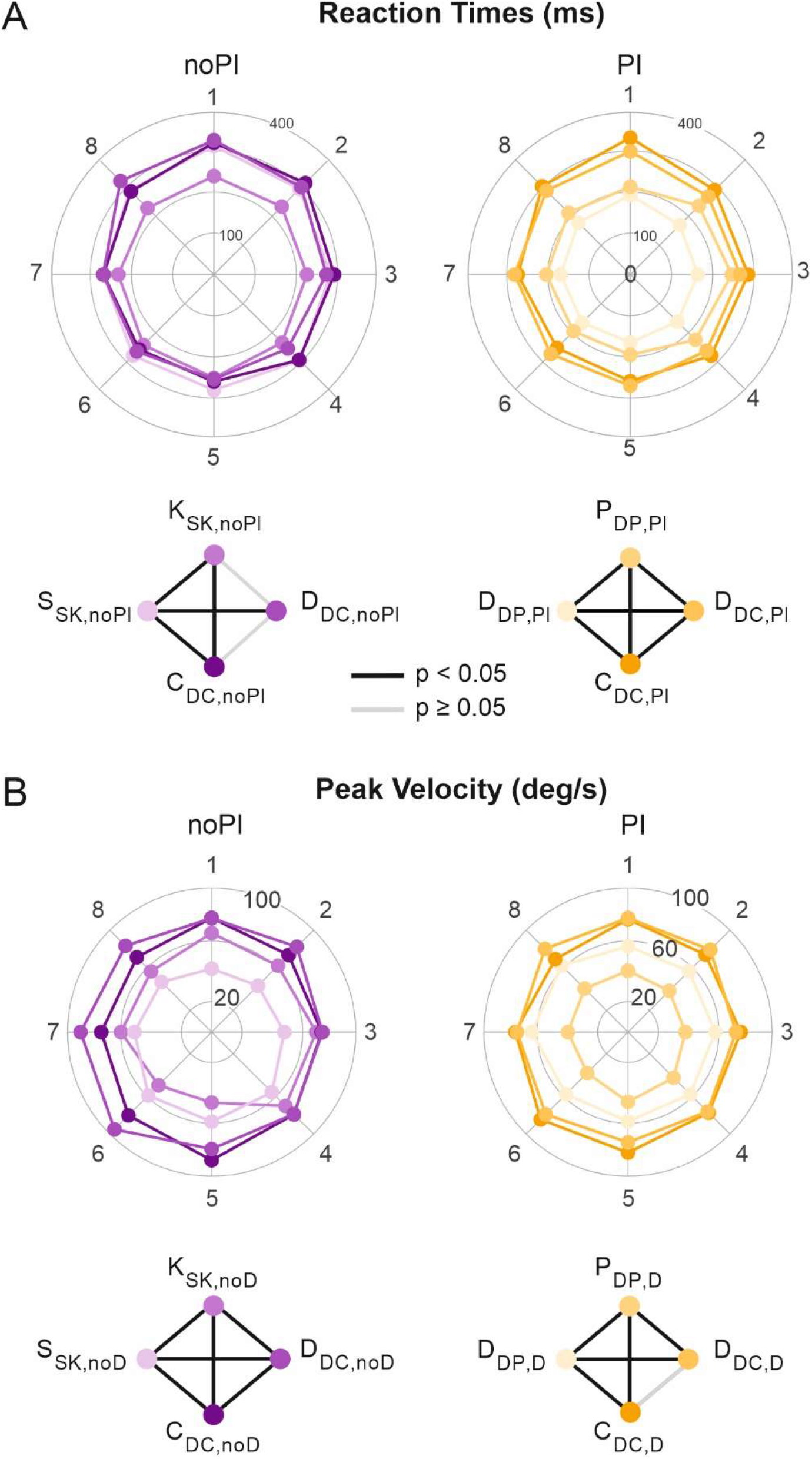
Inter-individual differences in kinematics during SOLO condition. Polar plots with median RTs (A) and median PVs (B) across directions measured for each monkey during the SOLO condition, for both noPI (purple) and PI groups (orange). Different color tints indicate different animals, as specified in the diamonds below each polar plot. Diamonds provide a graphical representation of statistical pairwise comparison in RTs (or PVs) between two monkeys. Black/gray segments refer to significant/non-significant differences (Dunn-Šidák’s test, p<0.05) between RTs (or PVs) of the animals reported at the vertices of the diamond. For each monkey it is reported the database of reference.

### Individual and inter-individual differences in TOGETHER performance

To test whether inter-individual differences, which emerged in SOLO condition, correlated with the dyadic performance during the TOGETHER trials (**Fig. 5**), we correlated the differences in RTs and PVs (IID_RT_, IID_PV_) with the inter-cursor distance error rate (ER_ICD_), taken as a measure of the goodness of joint performance. We assumed that this index reflects with more accuracy failures in inter-individual motor coordination. Interestingly, we found a positive and significant correlation between ER_ICD_ and the entity of both IID_RT_ and IID_PV_, only for PI sessions, that is when the type of action was pre-instructed in advance (**Fig. 5A**). There was no evidence instead of such correlation when the future action context was not pre-cued (noPI; **Fig. 5B**). This suggests that when the animals are prepared to act together (PI task) they modulate their action based on the kinematics of the partner, and the higher is the difference in behavior between coagents, the higher is the chance to fail. In other words, joint behavior is facilitated by the similarity of kinematics parameters, when animals are pre-instructed. On the contrary, when the monkeys are not prepared to act jointly (absence of a pre-cue) the error rates are not related to the inter-individual differences, as if the action of each animal is planned independently from the partner’s behavior.

**Fig. 5.**
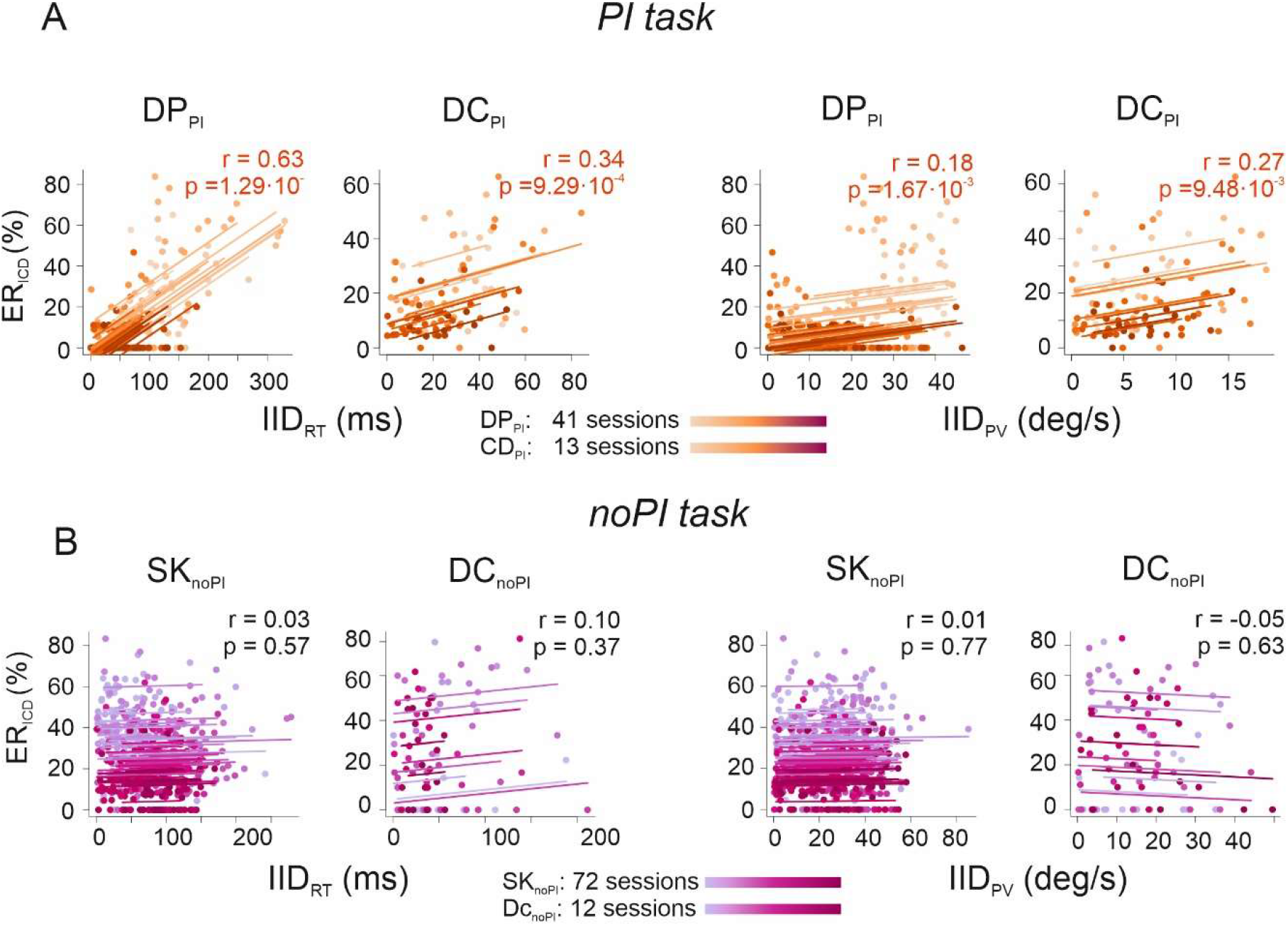
Correlations between inter-individual differences (IID) and joint-action performance. Repeated measure correlation between mean IID_RT_ and error rates (ER_IDT_) and between mean IID_PV_ and ER_IDT_ for PI (**A**) and for noPI PI (**B**) datasets. Each point represents the mean values of the two associated variables measured in one direction (1 to 8) in one session. Different sessions are color coded as reported in the legend.

### Effect of pre-instruction on monkeys’ kinematics

Given the substantial differences that emerged between the PI and noPI task, we analyzed which particular aspect was influenced by the presence of the pre-cue, finally leading to a facilitation of joint-action performance. To this purpose, we contrasted the RTs and PVs as measured in the two task conditions (noPI, PI) after pooling the behavioral data of all monkeys (**Fig. 6 A-B**) belonging respectively to the noPI and PI datasets. To exclude that the observed effects were associated to data pooling, as a control, the same analysis was repeated on the dataset of couple DC (Monkey D and C), which was tested in both versions of the task. The results showed that the pre-instruction about action condition reduced the RTs both in SOLO and TOGETHER trials (**Fig. 6A**), and this reduction was even higher for joint action trials respect to solo action. This result was confirmed for monkeys C and D.

**Fig. 6.**
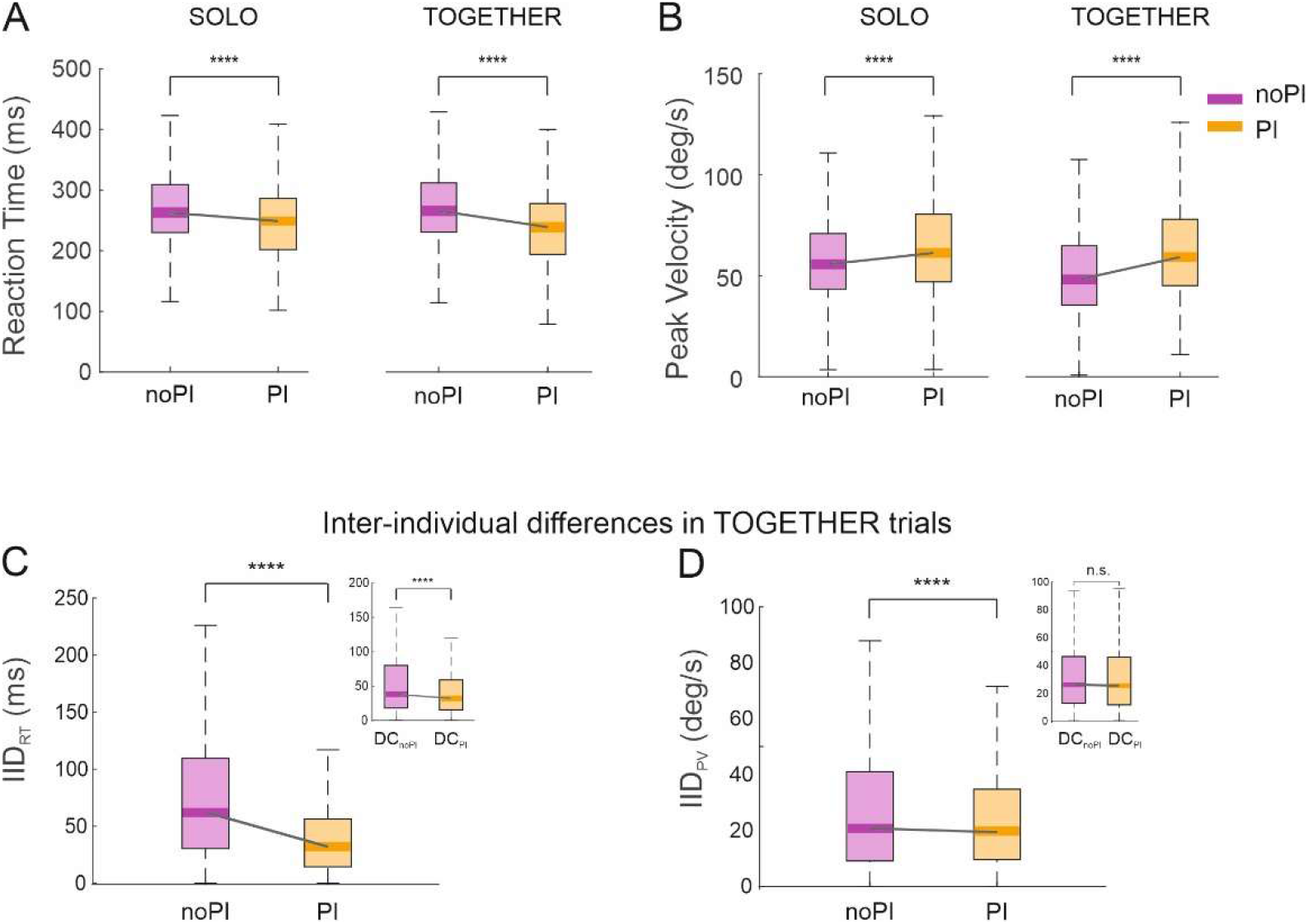
Influence of pre-instruction on movement condition on kinematics. **A-B**. Comparisons of RTs (**A**), or PVs (**B**), computed in noPI and PI task, during SOLO and TOGETHER trials. Boxplots refer to pooled data from all datasets. **C-D**. Comparison of inter-individual differences in RTs (**C**) or PVs (**D**), computed during TOGETHER trials, for the noPI vs PI datasets. Main plots refer to pooled datasets. In C and D top-right subplots represent data obtained from couple DC, which was tested in both noPI and PI task. **** = p < 0.0001 (Wilcoxon rank sum test).

As for PVs, a significant increase of this variable was observed in the pre-instructed trials, when pooling all datasets, both in SOLO and TOGETHER conditions (**Fig. 6B**), suggesting that overall, the animals were faster in guiding their cursors when they were pre-cued. However, this effect was not confirmed for monkey C and D, which either did not change (monkey D) or decreased (monkey C) their PVs. Therefore, the result obtained on PVs, by pooling all databases, could be due to intrinsic behavioral differences among subjects, in particular when comparing data from SK dataset tested in noPI task and the data obtained from couple DP tested in PI condition, rather than consequence of the type of task per se.

In summary, we observed that the presence of pre-instruction led to a systematic RTs reduction. Variable results were obtained instead for the execution phase, across animals. We postulate that pre-cuing action context, by inducing a significant RT reduction, might provide an optimal “kinematic setting” that ultimately could make inter-individual motor coordination easier.

### Pre-instruction optimizes inter-individual coordination in TOGETHER condition

Inter-individual differences measured in SOLO condition correlated with TOGETHER performance only for couples performing the PI task. We also found that pre-instruction modifies each monkey’s kinematics, particularly making action onset faster. Our hypothesis was therefore that this RT reduction, emerging thanks to an anticipatory formation of a dyadic plan, might facilitate the inter-individual motor coordination, by reducing behavioral inter-individual differences between the interacting animals. In other words, under this hypothesis the pre-instruction would provide an optimal “kinematic setting” that ultimately explains the higher success rate when a pre-cue is provided. To this aim in the TOGETHER condition we measured the inter-individual differences between the RTs and PVs of interacting monkeys, in the two task versions (noPI, PI) separately.

We found that, by pooling the different datasets, in the PI task the differences between the RTs (IIDRT) of the interacting monkeys (measured in TOGETHER trials) were significantly smaller, with respect to the noPI case (**Fig. 6C**; Wilcoxon rank sum test, All: z=36.12, p=1.22e-285). The same result was obtained on couple DC, which was exposed to both task versions. Interindividual differences in PV (IID_PV_), instead, despite the decrease observed when merging the data from different couples of the noPI and PI groups, did not change significantly for couple DC (**Fig. 6D**). Therefore, we cannot exclude that the apparent IID_PV_ reduction can be explained by task-independent differences between behavior of SK (tested in noPI) and DP (tested in PI).

In conclusion, our findings indicate that precueing indeed facilitates joint performance, by leading to a higher degree of coordination between individuals, intended as a greater similarity between the behavior of the two interacting animals. This similarity was particularly associated to a higher synchronization in the starting phase of joint action, as demonstrated by the significant reduction of the monkeys’ IID_RT_.

## DISCUSSION

The aim of this study was to analyze, in a quantitative fashion, the cognitive-motor processes guiding collective behavior in macaque monkeys, in particular when inter-individual motor coordination is required to successfully accomplish a shared task. These aspects are of special interest to understand the roots of advanced forms of cognitive-motor abilities of humans, such as when implementing complex inter-personal interactions and coordination.

### Evidence of “we-representation” in monkeys

There is a consolidated ethological knowledge that non-human primates, as well as other species, are able to cooperate when coordinated actions are mandatory to achieve a common goal. In natural environments, chimpanzees (Boesch & Boesch, 1989) show different level of cooperation during hunting. In particular, the third level of cooperation, that according to Boesch & Boesch’s occurs when two individuals display similar actions on the same goal and try to synchronize their behavior, has been confirmed also when apes are tested in more controlled experimental settings. The ability and attitude to coordinate individual actions with those of co-specifics have been shown often by using the bar-pull apparatus introduced by Crawford (1937) on chimpanzees, and subsequently on orangutans (Chalmeau et al. 1997) and capuchin monkeys (Mendres & deWaals, 2000).

In a highly controlled experimental setting, our previous studies have shown that macaques are a good model to study the neural and cognitive bases of the motor coordination required by a joint-action context. Macaques are in fact able to modulate their own action kinematics, as to adapt their own behavior (Visco-Comandini et al. 2015; Ferrari-Toniolo et al, 2019) to that of their co-acting partner. They are also able, when acting together, to apply behavioral strategies facilitating temporal synchronization and accuracy in spatial coordination.

The main goal of the present study was to explore the task contingencies favoring, therefore improving, monkeys’ performance in a joint action behavior. A way to approach this issue is to recourse to an “interactionism” perspective, holding that individuals, primed to interact, may count on an interpersonal awareness of their shared intention, rather than on continuous reciprocal mindreading by single agents (Gallotti & Frith, 2013). Theories of shared intentionality in humans claim that co-agents engaged in joint contexts represent their actions as pursued together based on a sense of ‘we-ness’, a fundamental pillar in psychology of collective behavior. Several human studies offer empirical support to the idea that joint action execution and planning involve a motor representation of collective goals (Novembre et al. 2012, 2014; Della Gatta et al. 2017; Kourtis et al. 2019; Sacheli et al. 2018), where an accurate Dyadic Motor Plan might subserve an efficient interaction. In this study we tested the existence of a ‘we-representation’ in monkeys, that, if present, might ameliorate the efficiency of interindividual coordination. To this aim we have designed two versions of an isometric joint action task, which differed by the presence/absence of a pre-instructing cue about the type of action (SOLO or TOGETHER) to be performed.

There is overwhelming evidence that movement pre-cueing about a future action influences motor planning (Rosenbaum, 1980). Pre-instructing about the effector to be used, movement direction, force, and movement extent (Bonnet & MacKay, 1987; Deiber et al, 2005; Jentzsch & Leuthold, 2002) lead to faster (i.e., shorter RTs) and more accurate responses, associated to an increased amount of advance information, which are likely to be processed at central level in motor programs. Our hypothesis was that, if the monkeys were able to access a ‘we-mode’ representation of the interactive scene at motor level, once the instruction to act together with a partner is provided in advance, this anticipatory representation should be beneficial to the execution of joint behavior, making it faster and more accurate, as much as any preliminary hint about motor features of the task to be performed is beneficial to its execution. On the contrary, once the relative pre-instruction is provided, the lack of such motor “we-representation” should have no effects on the motor response or on the goodness of interactive performance. In agreement with our previous results obtained both in humans and non-human primates tested in similar isometric tasks (Visco-Comandini et al. 2015; Satta et al. 2017), we found that the joint performance deteriorates individual animals’ behavior as compared to solo action. Interestingly, we found that when the type of action was cued in advance throughout a dedicated delay period, joint behavior improved significantly, as documented by the success rates (**Fig. 2**). Noticeably, the pre-instruction did not positively influence undistinguishably the performance of all trial types (i.e. SOLO and TOGETHER) but seemed to be beneficial particularly for dyadic behavior (**Fig. 2C**). The effect of pre-cueing was reflected also on the RTs. First, the anticipatory preparation to the future action-type led to a significant reduction of the RTs in the TOGETHER conditions with respect to SOLO action (**Fig. 3**). Second, this reduction has to be considered in the context of an overall decrease of RTs (observed in both SOLO and TOGETHER trials) associated to the presence of the pre-cue (**Fig. 6A**), as it will be discussed below.

These findings are in line, not only with the effects that advance information exert on motor planning but also with similar results obtained in humans tested specifically during joint behavior (Kourtis et al. 2019). This study showed that pre-cueing information about joint-action context affects our planning processes and facilitates dyadic performance, probably thanks to a predictive cognitive and sensorimotor “We-representations”, emerging at the group-level.

As a further evidence of this representation and coherently with human studies, we found that in monkeys the pre-instruction about future action type exerts beneficial effects on interindividual coordination. This is indicated by the significant decrease in the inter-individual differences between hand action onset times (reaction times) and force application speed (peak of velocity), typical of the TOGETHER trials (**Fig. 6C-D**). This suggests that pre-cueing might trigger a “we-mode” representation that leads to a significant amelioration of joint performance.

Intriguingly, a “we-representation” in primates can be inferred also from the ethological observations on the motivations driving the animals to pursue a joint behavior. A study on capuchins monkeys (Brosnan et al. 2006), using the bar-pull paradigm, showed that joint action was not affected by the equity of the rewards delivered to each of the two animals, but was rather positively influenced by the presence of high-value rewards offered to the dyad. Each animal was keen to pull the bar even when its partner in a given trial received a better offer. The joint success was therefore related to the reward value offered to the couple (“we” and not “I” from the monkeys’ perspective). Capuchins were instead sensitive to inequity, if systematically only one of the two partners regularly received the higher-value reward. This evokes the existence of a “we-mode” representation also in the processes which motivate the joint performance.

At neuronal level, we propose that this “we-representation” might have a correlate in the activity of premotor neurons recorded from the brains of macaques engaged in a joint action experiment identical to that of the present study (Ferrari-Toniolo et al 2019). These neurons, defined as “joint-action cells”, changed their firing when a given action was performed jointly with a partner, as compared to when the identical action was executed in a solo fashion. The same study has shown that their functional role during dyadic performance was putatively grounded on a predictive coding (see Friston, 2005) of own and other actions, not necessarily based on mirror mechanisms (Kilner et al. 2007), being only a subpopulation of the joint action cells characterized by mirror-like responses. This is in line with the predictive nature of the “we-representation” that facilitates the performance of dyadic behavior.

Neural markers related to representation of joint behavior have been shown in the EEG study by Kourtis et al (2019), which demonstrated that pre-specifying joint configuration modulate the amplitude of several EEG indices of cognitive (P600) and sensorimotor representations (alpha/mu rhythm and late CNV) related to action planning.

In conclusion, our results offer empirical evidence of a sensorimotor representation of collective behavior macaques. Pre-cueing favors the emergence of a We-mode which exerts an overall beneficial effect on the inter-individual coordination.

### An optimal ‘kinematic setting’ for acting together

We have shown that monkeys take into consideration the pre-cue about the action type, which lead the animals to initiate movement faster, and to increase their chance to coordinate successfully. The next question is to understand why the pre-instruction resulted in the improvement of joint action performance, and the origins of the beneficial effects.

First, from the comparison of the kinematics changes in a dyadic context with respect to SOLO actions, monkeys implemented an active inter-individual motor coordination strategy by anticipating the initiation of their movement (Visco-Comandini et al. 2015). It is known that speeding up action onset, result in less variable RTs both in humans (Repp, 2005; Wagenmakers & Brown, 2007) and in monkeys (Visco-Comandini et al 2015). This implies that RTs decrease facilitate inter-individual coordination, by reducing the variability of action onset time, which in turn increases behavioral predictability. Even if not intentionally, these phenomena might constitute a “coordination smoother” (Vesper et al. 2010), as observed in human studies (Vesper, et al., 2011; Masumoto & Inui, 2013; Sacheli et al., 2013). Interestingly, in our experiment RTs reduction is not only observed when comparing SOLO vs TOGETHER trials, as in our previous experiments (Visco-Comandini et al 2015; Ferrari-Toniolo et al. 2019), but the pre-instruction contributed a further RTs decrease, which was stronger in the joint-action condition. Overall, the RTs variations, which resulted in a reduction action planning variability, led to an increase in synchronicity in action initiation. Therefore, the presentation of a pre-cue promotes changes in kinematics known to favor inter-individual motor coordination. In particular, the tendency to anticipate the action initiation might foster the demonstrated ability of the dyad to decrease inter-individual differences while performing the task together. This aspect alone could favor motor coordination during joint trials, irrespective of other motor variables, such as direction and/or speed. It is worth noticing, that our PI version of the task did not prime the action’ goal, but only the action context (SOLO or TOGETHER). We conclude that prompting in advance the action context increased the chances of dyadic success, by establishing an optimal “kinematic setting” that ultimately facilitates inter-individual motor coordination.

Finally, in line with our previous observations, we confirmed that spontaneous behavioral invariances, generally associated to joint-action trials, are not always implemented by each agent. Such spontaneous strategies are sometime not sufficient to cope with the high demands of motor interactions, particularly when marked inter-individual differences, known to be associated with variability in brain structures (Kanai & Rees, 2011), exist between the two coagents. These diverse “individual motor signatures”, highlighted also in the present experiment (**Fig. 4**), might interfere with a smooth interpersonal coordination (Słowiński et al. 2016). We found that greater similarity signatures positively correlate with the success rates. Remarkably, this correlation was significant only when the couples performed the PI version of the task (**Fig. 5**). Provided that as humans (Schmidt et al., 2011; Sebanz & Knoblich, 2009, Sacheli et al., 2012), also macaques are able to modulate their motor behavior appropriately for a spatio-temporal coordination with their partners (Visco-Comandini et al 2015; Ferrari-Toniolo et al. 2019), our interpretation is that in the PI task, contrary to the noPI one, the animals could benefit of the pre-cue to better prepare tacking the potential differences in the kinematics of their mates relative to their own behavior.

## Author contributions

ABM, SFT and RC designed the study; AS, LB, LEN, RG, SFT, RC and ABM performed the experiments; IL, LB and ABM analyzed the data; IL, ABM wrote the paper.

## ACKNOWLEDGEMENT

We are grateful to Giacomo Novembre for commenting on the manuscript, and to Stefano Ferraina for his valuable help during experimental procedures. We acknowledge the support of the Istituto Italiano di Tecnologia to RC. This work was supported by the MIUR of Italy (PRIN 2017 to ABM, Grant n. 201794KEER_002).

